# scRMD: Imputation for single cell RNA-seq data via robust matrix decomposition

**DOI:** 10.1101/459404

**Authors:** Chong Chen, Changjing Wu, Linjie Wu, Yishu Wang, Minghua Deng, Ruibin Xi

## Abstract

**Motivation:** Single cell RNA-sequencing (scRNA-seq) technology enables whole transcriptome profiling at single cell resolution and holds great promises in many biological and medical applications. Nevertheless, scRNA-seq often fails to capture expressed genes, leading to the prominent dropout problem. These dropouts cause many problems in down-stream analysis, such as significant noise increase, power loss in differential expression analysis and obscuring of gene-to-gene or cell-to-cell relationship. Imputation of these dropout values thus becomes an essential step in scRNA-seq data analysis.

**Results:** In this paper, we model the dropout imputation problem as robust matrix decomposition. This model has minimal assumptions and allows us to develop a computational efficient imputation method scRMD. Extensive data analysis shows that scRMD can accurately recover the dropout values and help to improve downstream analysis such as differential expression analysis and clustering analysis.

## 1 Introduction

Single cell RNA sequencing (scRNA-seq) technology can profile whole transcriptome at single cell level and has great potential for biological and medical applications(Tang *et al.*, 2009; Islam *et al.*, 2011; Hashimshony *et al.*, 2012; Ramsköld *et al.*, 2012). Many previously difficult questions can now be addressed with scRNA-seq, such as investigation of cellular heterogeneity, characterization of early embryonic cells and circulating tumor cells(Ting *et al.*, 2014; Kharchenko *et al.*, 2014; Moignard *et al.*, 2015; Deng *et al.*, 2014). After reverse transcription, current scRNA-seq technology usually requires heavy amplification owing to the picogram level of RNAs in a single cell. During the reverse transcription and the amplification step, RNA transcripts might be missed and consequently not detected in the following sequencing. This is called the dropout problem. In addition, since single cells only express a proportion of genes, many zeroes in scRNA-seq data are true zero expressions. As a result, scRNA-seq data consists of two types of zeroes, truly unexpressed zeroes and dropout zeroes.

Researchers have developed scRNA-seq data analysis methods from several perspectives. Most of these methods are designed for specific statistical analysis such as clustering, dimension reduction(Kiselev *et al.*, 2016; Lin *et al.*, 2017; Wang *et al.*, 2017; Bacher *et al.*, 2017; Pierson and Yau, 2015), and differential expression (DE) analysis (Kharchenko *et al.*, 2014; Vallejos *et al.*, 2016; Jia *et al.*, 2017). Another class of methods (Van Dijk *et al.*, 2017; Li and Li, 2018) focuses on imputing the dropout values. After proper imputation, available methods for bulk RNA-seq data may be directly applied to the imputed data and many different types of statistical analyses, such as differential expression analysis, clustering and dimension reduction can be directly performed. MAGIC(Van Dijk *et al.*, 2017) is a diffusion based imputation method. Roughly speaking, MAGIC performs a “soft” clustering after constructing a Markov transition matrix and replaces a gene’s raw expression with its weighted mean expression in a cluster, which greatly reduces the noise in the data. Although the noise reduction may be beneficial for a few subsequent analyses such as visualization, the large alteration of the original data can also be misleading. The method scImpute(Li and Li, 2018) imputes gene expression by inferring an expression relationship among single cells based on non-dropout genes and applying this relationship to dropout genes. An implicit assumption of scImpute is that the relationship of a dropout gene’s expression in one cell with this gene’s expression in other cells can be inferred from expression data of non-dropout genes. This assumption is quite strong and may not hold for many scRNA-seq data.

In this paper, we develop a single cell RNA-seq imputation method scRMD based on the robust matrix decomposition. scRNA-seq data often contain single cells from a few different, possibly unknown, cell types. Single cells of the same type should have similar expression pattern and thus it is reasonable to assume that the underlying true expression matrix is approximately a low rank matrix. In addition, the dropout events should be relative sparse in the expression matrix. Based on these two assumptions, we propose to impute the scRNA-seq by minimizing an objective function containing a low rank and a sparsity penalty term. An efficient alternating direction method of multiplier (ADMM) is developed to minimize the objective function. Extensive data analyses show that, compared with other methods, scRMD gives more accurate imputation and can improve subsequent clustering analysis and differential expression analyses.

## 2 MATERIAL AND METHODS

Given n single cell RNA-seq data with p genes, let *y_ij_ ≥* 0 be the normalized expression (Supplementary Text) of the *i*th gene in the *j*th cell. If there were no dropout, the normalized expression *y_ij_* would be the true expression *x_ij_* plus a random measurement error *e_ij_*. However, if dropout occurs, the measured expression *y_ij_* would just be the measurement error *e_ij_*. Denote *s_ij_* = *x_ij_* if the *i*th gene of the *j*th cell is dropped out and *s_ij_* = 0 if otherwise. Then, we have *y_ij_* = *x_ij_* − *s_ij_* + *e_ij_* for any *i* = 1,*…*, *p* and *j* = 1,*…*, *n*. In matrix form, we have

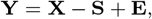

where ***Y*** = (*y_ij_*)_*p*×*n*_, ***S*** = (*s_ij_*)_*p*×*n*_and ***X*** = (*x_ij_*)_*p*×*n*_.

The dropout events should not occur too often (compared with the size of the expression matrix), and we can reasonably assume that S is sparse. In addition, cells can usually be clustered into a few cell types. Cells within the same cluster tend to share similar expression patterns. Specifically, let *l_i_*_,*c*_be the mean gene expression level of the *i*th gene of the cells in type *c*. Suppose that cell *j* belongs to cell type *c*(*j*). The *j*th cell’s *i*th gene expression can be written as *x_ij_* = *l_i_*,*c*(*j*) + *f_ij_*, where *f_ij_* is the *i*th gene’s random deviation from its mean expression level in the *j*th cell. Thus, we have ***X*** = ***L***+***F***, with ***L*** = (*l_i_*,*c*(*j*))_*p×n*_and ***F*** = (*f_ij_*)_*p*×*n*_. Since the number of cell types is usually much smaller than the number of cells, the rank of ***L*** should be much smaller than n. Consequently, the true expression ***X*** exhibits an approximate low rank structure in the sense that there are only a few large singular values followed by many weak ones for the expression matrix (Figure S1A). In summary, the gene expression matrix ***Y*** can be decomposed as

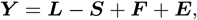

where ***L*** is a low rank matrix, ***S*** is a sparse matrix, and ***E***, ***F*** represent the measurement errors and random fluctuations of the cells around their mean expression levels. Since both ***E*** and ***F*** are random, we write ***E***_1_ = ***F*** +***E*** and thus ***Y*** = ***L****−****S***+***E***_1_. Figure 1B demonstrates an illustrative example for such decomposition. In statistics and machine learning literature, such decomposition is called the robust matrix decomposition (RMD)(Hsu *et al.*, 2011). Therefore, we may estimate ***L*** and ***S*** by minimizing

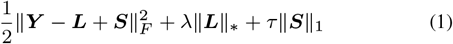

subject to ***L***, ***S*** *≥* 0 (the inequalities should be understood elementwisely), where ||⋅||*F*,||⋅||_*_, and *||* ||_1_ are the Frobenious norm, nuclear norm, and elementwise *ℓ*_1_ norm of a matrix. The nuclear norm penalty is to encourage the low-rankness of ***L*** and the *ℓ*_1_ norm penalty is to encourage the sparsity of ***S***. The tuning parameters *λ* and *τ* in (1) control the strengths of regularization.

**Fig. 1.**
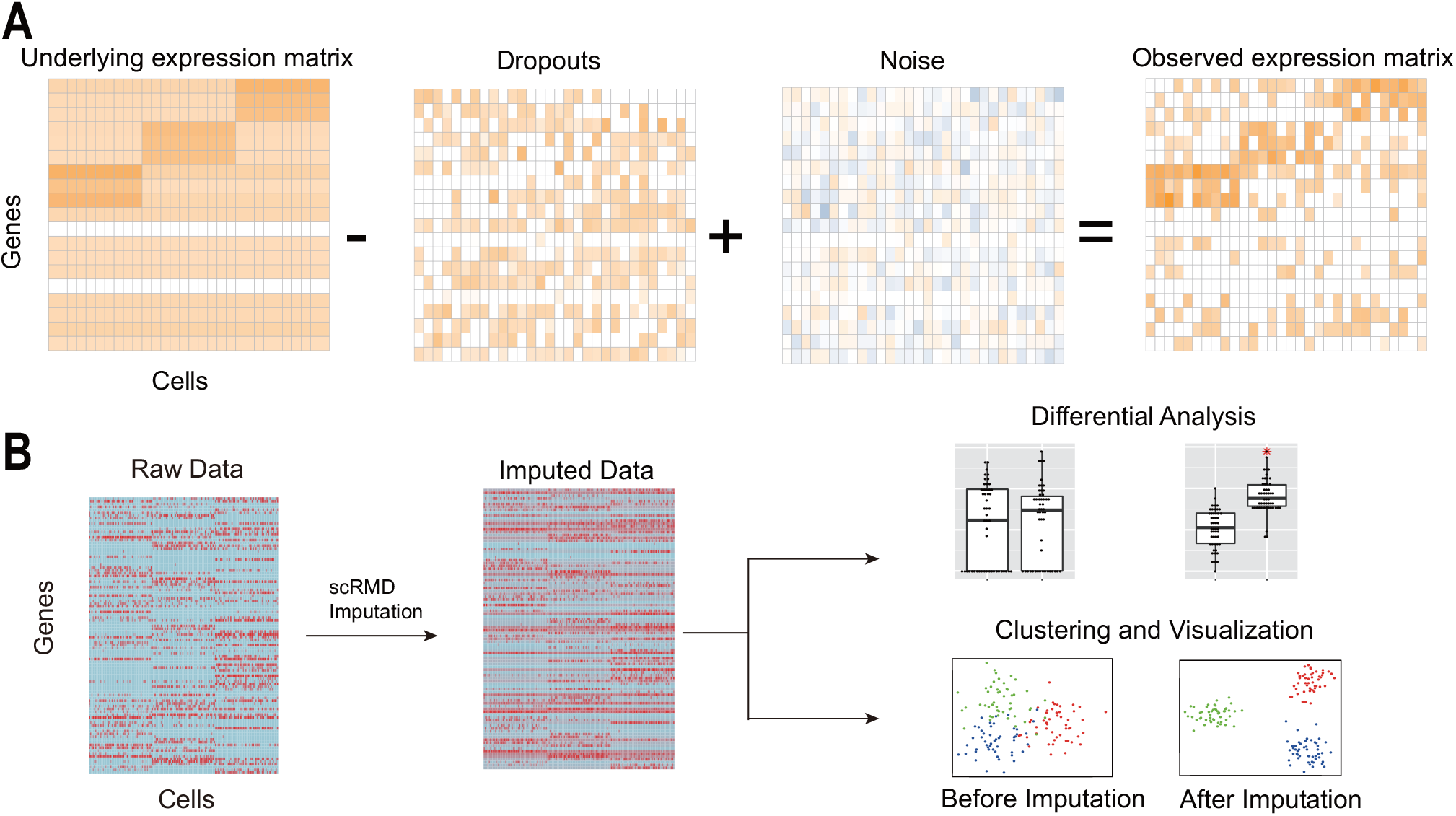
The outline of scRMD. (A) The observed expression matrix can be decomposed as the underlying low rank matrix **L**, the dropout matrix **S** and the noise matrix **E**1. (B) After scRMD imputation, available RNA-seq analysis methods can be applied to the imputed data.

For single cell sequencing data, we know that if the normalized expression *y_ij_* is large enough, the *i*th gene of the *j*th cell cannot have dropout and thus we must have *s_ij_* = 0. To incorporate such information into our method, when minimizing (1), instead of allowing *s_ij_* = 0 for all *i* and *j*, we will restrict *s_ij_* = 0 for all *i* and *j* with *y_ij_* > 0, where *c* ≥ 0 is a constant. More specifically, let Ω = {(*i*, *j*) : *y_ij_* ≤ *c*, 1 ≤ *i* ≤ *n*, 1 ≤ *j* ≤ *p*} and thus Ω^*c*^= {(*i*, *j*) : *y_ij_* > *c*, 1 ≤ *i* ≤ *n*, 1 ≤ *j* ≤ *p*}. The index set Ω is the candidate dropout event index set. Thus, we require *P*_Ω_(***S***) ≥ 0 and 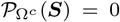 when minimizing (1), where *P*_Ω_(***S***) = (*P*_Ω_(*s_ij_*))_*p*×*n*_, and similarly *P*_Ω_*c* (***s***), is given by

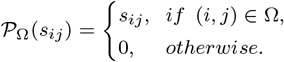

Hence, for a given candidate dropout set Ω, we propose to estimate ***L*** and ***S*** as the solution to the following optimization problem

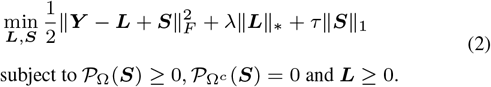

We develop an efficient ADMM algorithm for solving (2). For the tuning parameters, we use a random-matrix-based technology to choose *λ* and *τ* (Supplementary Text) and set *c* as the 0.05 quantile of the nonzero expression. Let ***L̂*** and ***Ŝ*** be the solution to (2). We use ***Y*** + *P_Ŝ>_*_0_(***L̂***) as the imputed data for downstream analyses. Note that ***L̂*** is the estimate of the group means. Hence, we just impute entries in the candidate dropout event set Ω as their group means. Another natural choice is to use ***Y*** +***Ŝ*** as the imputation data. Nevertheless, the elements of ***S***̂ are subject to biases due to the *ℓ*_1_ penalty(Van de Geer *et al.*, 2014) and ***Y*** + ***Ŝ*** would also subject to biases.

## 3 RESULTS

### 3.1 Simulation Analysis

We first perform simulation analysis to compare scRMD with scImpute and MAGIC. We consider two simulation setups. One is the DE analysis setup and the other is the clustering analysis setup. With these simulation data, we also compare the imputation accuracy of different algorithms. We set the total number of genes as *p* = 10, 100, and generate two groups of cells for DE analysis and three groups for clustering analysis. Each group has 50 cells. To generate the mean expression levels *μ_ik_* for the gene *i* in the group *k*, we first generate a base line expression *μ_i_* independently from the normal distribution *N* (1.8, 0.5^2^) for *i* = 1, …, *p*. We further randomly choose *ζp* (0 < *ζ* < 1) genes and set the corresponding *μ_i_* = 0. This allows us to generate true zero-expressed genes. For DE analysis, the first and the second group’s mean expressions are set as *μ_ik_* = *c_ik_μ_i_* (*k* = 1, 2). We set *c_i_*_1_ = 1.5 for the first 50 genes in the first group and *c_i_*_2_ = 1.5 for the next 50 genes in the second group. All other *c_ik_*’s are set as 1. For clustering analysis, mean values of the first 100 genes for each group are directly generated from the normal distribution *N* (1.8, 0.5^2^).

Given a cell in the *k*th group, the normalized expression *x_i_* of its *i*th gene is generated from a normal distribution 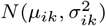, where *μ_ik_* is the mean expression of the *i*th gene in the *k*th cell group and *σ_ik_* = *μ_ik_σ_hetero_* + *σ_homo_*. Note that if *σ_hetero_* = 0, all genes will have the same gene expression variance; Otherwise, different genes have different variances. We vary *σ_hetero_* and *σ_homo_*to simulate different noise setups. If *x_i_* is generated to be less than 0, we set it as 0. Thus, *x_i_* is the gene expression without dropouts. To generate expression with dropouts, we generate a random number from the uniform distribution *U* (0, 1). If this random number is less than 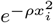, we set the observed expression *y_i_* = 0; Otherwise, we set *y_i_* = *x_i_*. The parameter *ρ* controls the dropout rate and we also vary it to compare the algorithms under different dropout rates. To account for technical noises, we further add a truncated normal noise *∊* = max{*N* (0, 1) *−* 1.5, 0} to the observed expression *y_i_*.

In the simulation, we set *ζ* = 0.3, 0.4, 0.5, 0.6, 0.7 and *ρ* = 0.1, 0.2, 0.3, 0.4, 0.5 to control different true zero rates and dropout zero rates. We also vary *σ_hetero_* = 0, 0.05, 0.1, 0.15, 0.2 and *σ_homo_* = 0, 0.1, 0.2, 0.3, 0.4 to account for different heterogeneous and homogeneous errors. When varying one parameter, all other parameters are fixed at the median of all possible values. For comparison, we also include the “oracle” method that uses the true expression matrix directly in downstream analyses. Under each setting, we repeat the simulation 50 times. For DE analysis, DE genes are identified by limma(Ritchie *et al.*, 2015) with Benjamini-Hochberg (Benjamini and Hochberg, 1995) adjusted p-values smaller than 0.05 and F-score are calculated to evaluate the accuracy. The mean imputation errors, defined as the mean absolute difference between the imputed and the true expression on the true dropout positions, are calculated for all methods. For clustering analysis, we perform dimension reduction by PCA and cluster the cells based on the first two principle components by the K-means algorithm. The number of clusters in the K-means algorithm is set as 3. The clustering results are evaluated by Adjusted Rand index (ARI) (Hubert and Arabie, 1985). All imputation methods are implemented in R version 3.4.0.

Figure 2A illustrates the F-scores of different methods for DE analysis. As expected, the Oracle method has the best performance since there is no dropouts while our scRMD achieves comparable results and outperforms other methods uniformly to a large margin. Since MAGIC tends to change all genes expression to their cluster means and significantly reduces the variances, DE analysis based on MAGIC-imputed data tends to give many false significant DE genes and hence the F-score of MAGIC is very low compared with other methods. Figure 2B shows the mean imputation errors of different methods based on the DE-analysis data. Overall, scRMD achieves the most accurate imputation results compared with other algorithms. However, when the dropout rate parameter *ρ* is small, scImpute have smaller mean loss than scRMD. This is because many large values are dropped out when *ρ* is small and scImpute prefers to impute missing values with relative large values. The ARI Figure 2C shows the ARI of the clustering analysis (see Figure S2 for an example of these clustering results). Similar to the DE analysis, Oracle is the best method as expected, while scRMD achieves comparable results in most cases and is better than scImpute, MAGIC and Raw. scImpute is sensitive to all parameters and only has good performance in cases with low zero rates and low noise levels. In summary, simulation studies show that scRMD is an effective imputation method that could lead to better downstream statistical analysis in single cell RNA-seq data. We finally report the mean computational time for imputing one simulation data, scRMD (3.6 ± 1s), scImpute (36.7 ± 0.2s), and MAGIC (0.3 ± 0.0s). MAGIC is the fastest method since it only involves matrix multiplication. In the meanwhile, scRMD only takes a few seconds and is about ten times faster than scImpute.

**Fig. 2.**
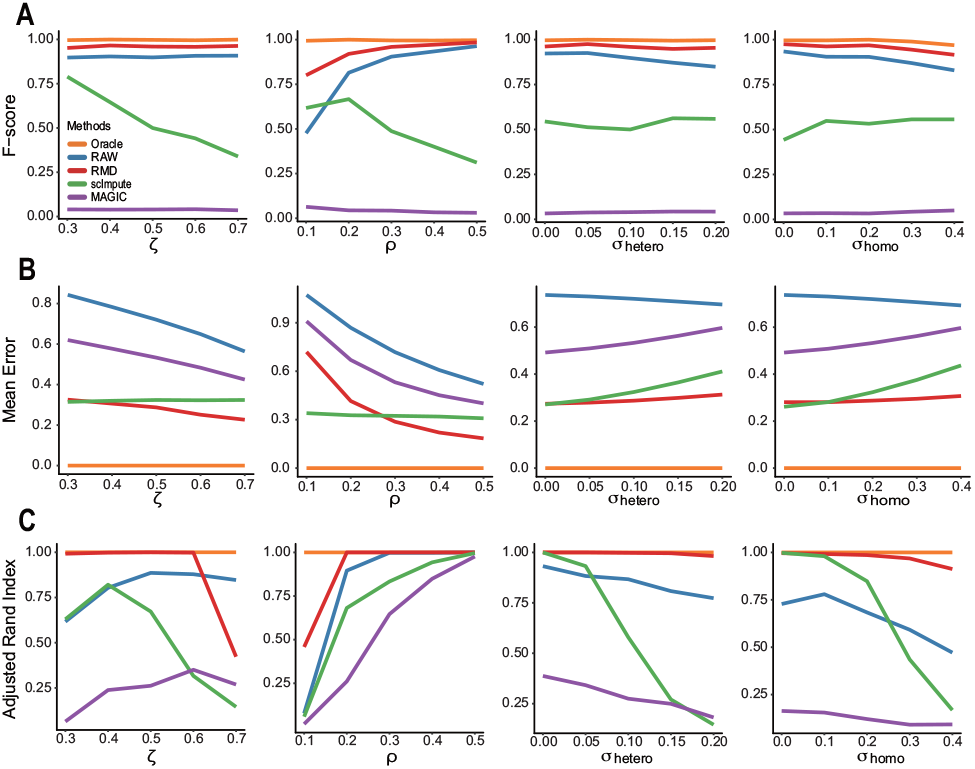
Comparison of algorithms by simulation. (A) The F-scores of the algorithms for DE analysis; (B) The mean imputation errors, (C) The adjusted Rand index of the clustering results based on imputation data given by different algorithms.

### 3.2 Imputation Accuracy in Real data

We evaluate the accuracy of the imputed data given by different algorithms using the Ziegenhain scRNA-seq data(Ziegenhain *et al.*, 2017). The Ziegenhain data contains 583 cells that were sequenced by six different methods, and five of them contain spike-in synthetic RNA transcripts with known concentration designed by the External RNA Control Consortium (ERCC)(Jiang *et al.*, 2011). Since we only know the relative expression level of these ERCC genes, we use the Pearson correlation of these ERCC genes between the imputed data and the known concentration level to study the imputation accuracy. The original counts or UMI data are first pre-processed (Supplementary Text). All data are imputed by scRMD, scImpute and MAGIC. Figure 3 shows the boxplots of the Pearson correlations between the imputed value and the known log concentration level. Compared with the raw data, all imputation methods improve the correlations. MAGIC seems the best method in terms of improving the correlation. However, we see that the correlations given by MAGIC have very small variances among single cells in all five data sets. This is again because MAGIC tends to replace genes’ observed expressions by their cluster means. scRMD gives larger, often significantly larger, correlation than scImpute in four of five sequencing data sets (Table S1). For example, in the SCRBseq data, the median correlation of scRMD is over 0.910, but that of scImpute is only 0.887.

**Fig. 3.**
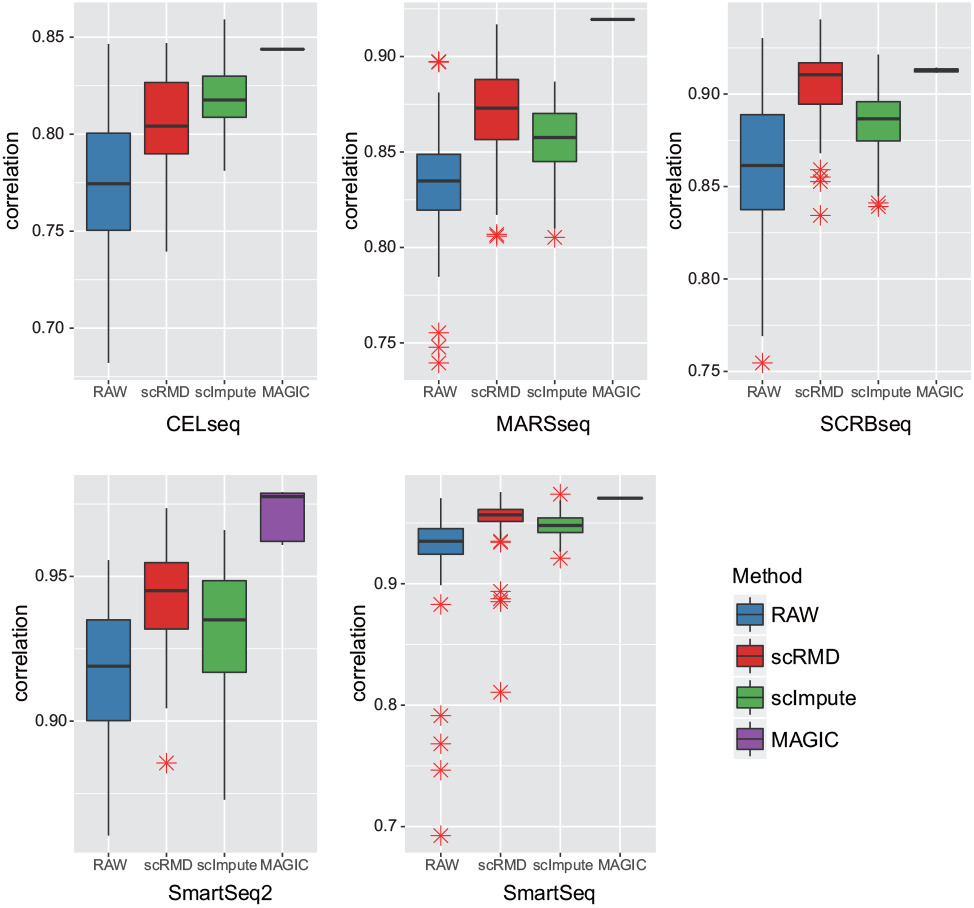
Pearson correlations between imputed data and known concentration of ERCC genes.

### 3.3 scRMD improves differential expression analysis

In this section, we show the effect of dropout imputation in terms of DE analysis in real data. Here, we consider two real data sets. One is the human embryonic stem cell data that consists of 212 H1 human embryonic stem cells (ESC) and 138 definitive endoderm cells (DEC) (Chu data)(Chu *et al.*, 2016). Another one is the hematopoietic stem cell data from young or old mice (Kowalczyk data)(Kowalczyk *et al.*, 2015). After removing very low coverage samples (total read count less than 10,000), the Kowalczyk data has 89 and 135 cells for the young and old mice, respectively. After imputation, we use limma to detect DE genes. Parameters of all algorithms are set as the default values.

The Chu data has 6 bulk RNA-seq data and we use these bulk data as the golden standard. Under the false discovery rate of 0.01 and a minimum fold-change of 2, limma identified 4,157 DE genes from the bulk data. We choose the top 100, 200, 500 and 1,000 genes ranked by p-values from the bulk data as the gold standard, which correspond to different level of stringency of setting the golden standard. We rank the genes detected in single cell data increasingly by their p-values. Then, we select the top k DE genes and calculate the overlap with golden standard gene sets. Figure 4A and S3A shows the overlap curves for a variety of k. The ratio between the overlap and the number of detected genes in single cell data is a measure of true discovery rate and the ratio between the overlap and the number of genes in the golden standard set is roughly the sensitivity of the DE detection. We see that scRMD uniformly outperforms other imputation approaches as well as the Raw method, especially if we set the golden standard set to be more stringent. The performance of scImpute is also good, while MAGIC cannot improve DE analysis over the raw data method. The above overlapping analysis with the golden standard set depends on how we choose the golden standard. To avoid the potentially biased performance evaluation due to such dependence, we further compare the ranks of the detected DE genes by each algorithm in the bulk data. More specifically, for each method, we take the top k DE genes and calculate the mean rank of these genes in the bulk data. Figure 4B clearly shows that the mean rank of genes given by scRMD is smaller than other algorithms, meaning that the DE genes detected by scRMD is more concordant with the bulk data. To further investigate effect of imputation on DE genes with high dropout rates, we select DE genes from the bulk data with more than 80% zeroes in single cell data and plot the p-values (Wilcoxon’s rank sum test(Wilcoxon, 1945) and limma) based on the imputation data against those based on the raw data (Figure 4C and Figure S3B, top panel). As a comparison, we also plot the p-values for top 500 non-DE genes ranked by p-values (Figure 4C and Figure S3B, bottom panel). For DE genes, all imputation methods generally give smaller p-values than the raw data and scRMD tends to give smaller p-values than scImpute, meaning than scRMD would be more sensitive in detecting these DE genes. MAGIC gives very significant p-values for both DE genes and non-DE genes, again shows that the imputation data given by MAGIC is not suitable for DE analysis. As a comparison, for non-DE genes, the p-values of scRMD and scImpute roughly lie on the diagonal line.

**Fig. 4.**
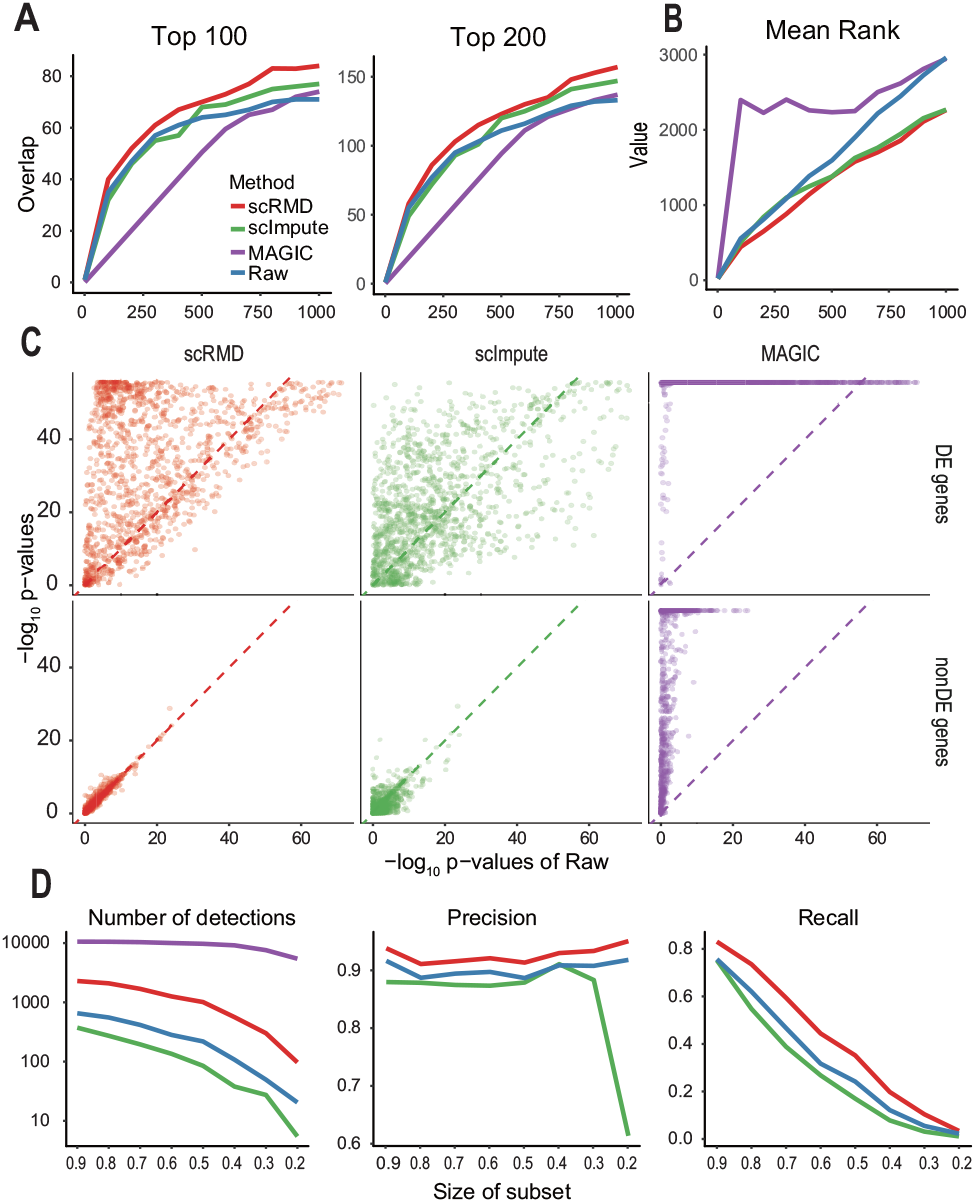
DE analysis in real data analysis. (A) The overlap of single cell DE genes with the golden standard gene sets. The golden standard gene sets are chosen as top 100 and 200 DE genes in the bulk data for the left and right panel, respectively. (B) Mean ranks in the bulk data of the detected DE genes by each algorithm. (C) The scatter plots of DE p-values based on the imputation data against p-values based on the raw data. Top panel corresponds to DE genes in the bulk data with more than 80% zeroes in the single cell data and the bottom panel is the non-DE genes in the bulk data. (D)The number of detected DE genes, the precision and the recall of each algorithm as a function of the proportion of cells used.

The Kowalczyk data does not have additional information that can be used as golden standard and we use down-sampling strategies to study the reproducibility of different algorithms. Under a false discovery rate of 0.05, based on the entire data set, the Raw, scRMD, scImpute and MAGIC detected 788, 2,577, 433, and 10,640 genes, respectively. We consider two types of down-sampling strategies here. The first one is to sample a subset of cells per group. We vary the subsample percentage from 20% to 90%. The first panel in Figure 4D shows the number of DE genes detected by each algorithm on different subsampled data. As expected, numbers of detections for all methods decrease as we decrease the numbers of cells per group. However, MAGIC almost constantly detect all genes as DE genes and hence are excluded in other comparisons. For each algorithm, we then use the DE genes detected based on the entire data as the “gold standard”(hence Raw, scRMD and scImpute have 788, 2,577, and 433 DE genes respectively) and calculate the precision and the recall on the subsampled data (Figure 4D, the second and third panel). All methods have relatively high precision regardless of cell number in the data, while the recall decreases when the cell number becomes smaller. We see that scRMD outperforms other algorithms uniformly, implying that the imputed data by scRMD is more stable and reproducible. The second down-sampling method is to decrease the library size for each cell by subsampling read counts from the raw count matrix. We subsample 10% of the original data, which leads to a significant decrease of nonzero proportion from 68% to about 22%. In this experiment we use the top 50, 100, 200, and 400 DE genes detected by each algorithm based on the raw data set as their own golden standard. Note that since scImpute can only detect 433 DE genes with entire data set, we set the largest number of golden standard DE genes as 400. Figure S3C shows the receiver-operating curves (ROC) for each algorithm when we vary the number of detections of each algorithm. We see that in general the area under curve (AUC) of scRMD is much larger than the other two algorithms and roughly similar to the RAW method.

### 3.4 scRMD Improves Clustering Analysis

We next investigate the performance of the imputation methods for clustering analysis. We consider four different data sets, the Usoskin data (Usoskin *et al.*, 2015) consisting of 622 cells from 4 different cell types, the Pollen data (Pollen *et al.*, 2014) having 249 cells from 11 cell types, the Ting data (Ting *et al.*, 2014) with 149 cells from 7 cell types and the Deng data (Deng *et al.*, 2014) with 268 cells from 10 cell types. After data pre-processing and imputation, we perform PCA and tSNE analysis (Maaten and Hinton, 2008) to reduce the data dimension. K-means algorithm is then applied to cluster cells, where K is chosen as the known number of cell types. We set the number of principle components as the number of cell types. The dimension for tSNE is set as 2. We also consider two clustering methods CIDR (Lin *et al.*, 2017) and SIMLR (Wang *et al.*, 2017), which were developed for clustering single-cell RNA-seq data. In addition, we also perturb the raw data and evaluate the performance of the imputation methods for clustering analysis under various perturbations. The first perturbation is to randomly sample 10% of the original data to evaluate the performance of the algorithms for low coverage data. In the second case, we randomly dropout gene expressions under the exponential decay model with a dropout probability 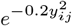. These two sampling strategies are repeated 50 times to account for randomness in sampling. In the third case, we only use genes whose mean expressions across cells are the lowest 25% among all genes. Since low-expressed genes are more likely to be affected by dropout events, this third perturbation also allows us to investigate the performance of the algorithms when there is a higher dropout rate in a model free manner. We call these three perturbations, the down-sampling perturbation, the model-based-dropout perturbation and the low-expression-gene perturbation, respectively. The mean zero rates under different perturbations are shown in Table S1. We see that the zero rates are significantly increased under different perturbations.

Figure 5A-B shows the clustering results of the Usoskin and Ting data (See Supplementary Figure S4C-D for the Pollen and Deng data). For the full data clustering analysis, the improvement of clustering by scRMD over the raw data method varies depending on the data. The improvement by scRMD is minimal for the Pollen and Deng data, but the improvement is essential for the Ting and Usoskin data. A likely reason is that the Ting and Usokin data have higher proportions of zeroes and hence higher dropout rates than the other two data sets (Table S2). It is clear that scRMD performs better than other imputation methods in most cases. Interestingly, in many cases, scRMD performs similarly to scImpute and MAGIC on the full data set (except the Usoskin data), but scRMD almost always performs much better than the other imputation methods for the perturbed data sets. This shows that scRMD can give more accurate imputation than other methods when there are more dropouts. For the Usoskin data, scRMD performs much better than the other imputation methods even on the full data set, probably because the Usoskin data has much higher zero rate (78%) than other data sets (Table S2). Another reason that the performance of scImpute and MAGIC is not satisfactory is that scImpute tends to impute large values to the dropout entries (Figure S4A), and MAGIC tends to change both zero and non-zero entries that leads to a huge loss of original information. CIDR also does imputation before clustering, but we can see that the performances are significantly better on imputed data by scRMD in most cases, which implies scRMD gives better imputation results.

**Fig. 5.**
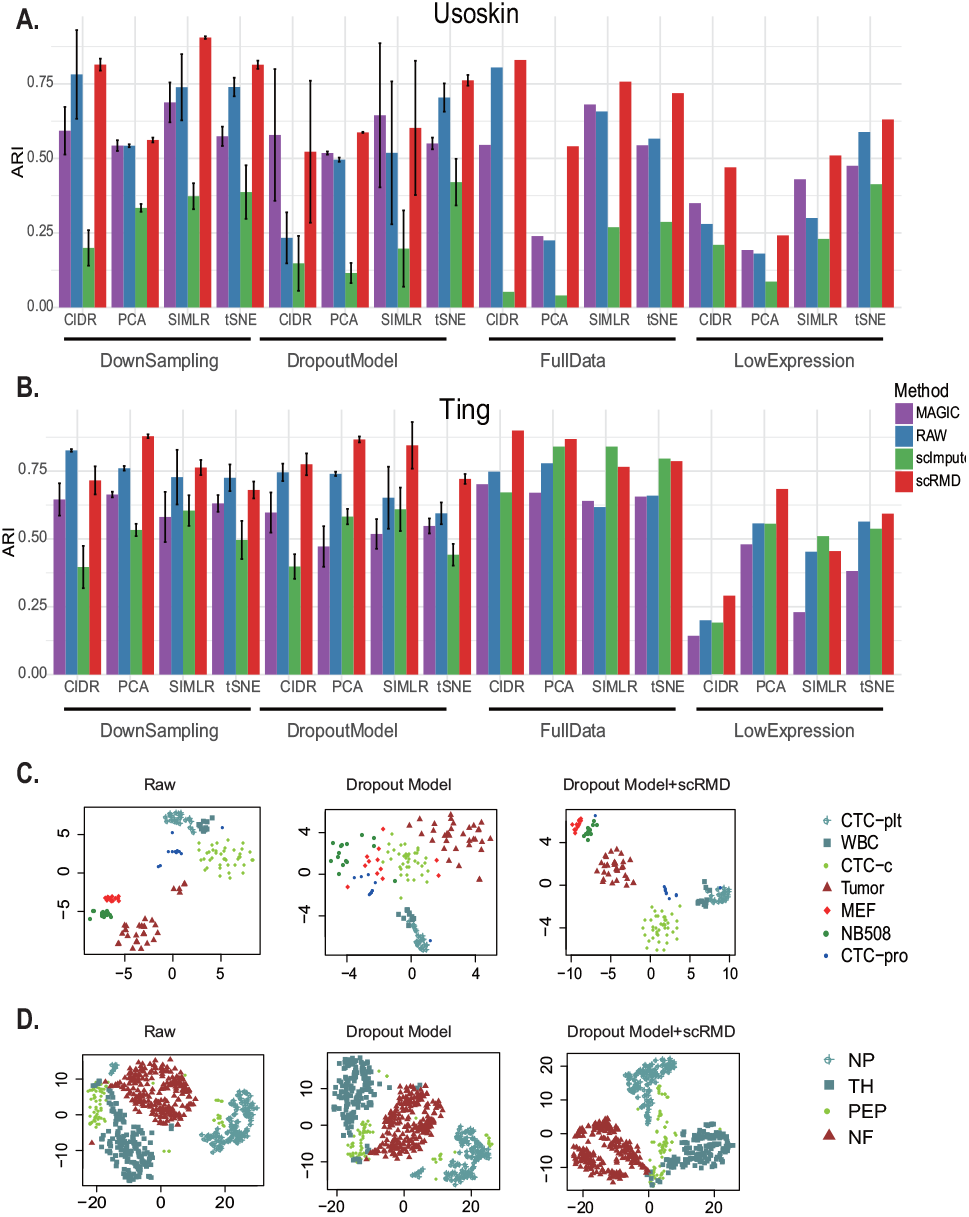
Clustering on real data. (A,B) The ARIs of the clustering results for the full data and the perturbation data for the Usoskin data and the Ting data, respectively. (C) For the Ting data set, the two dimensional tSNE scatter plot of the raw data, one of the model-based dropout data and the scRMD imputation imputed model-based dropout data. (D) Similar to (C), but for the Usoskin data.

To further illustrate the improvement of clustering by scRMD, we plot the two dimensional tSNE results for the Ting data and the Usokin data in Figure 5C-D, where the left panels show the raw data, the middle panels one of the model-based-dropout perturbation data and the right panels the scRMD imputation of the dropout perturbation data. For the Ting raw data, cells in the same cell type largely cluster together and cells from different cell types are further away from each other, but four cells of the Tumor type are separated from the majority of the Tumor cells. After we introduce more dropouts by the dropout perturbation, different cell types can no longer be as well separated as the original raw data. However, after we perform the scRMD imputation, different cell types can again be clearly separated. The cells of the same type are even more tightly clustered. For example, unlike in the raw data plot, the Tumor type cells are now all closely clustered together. Similar phenomenon is also observed in the Usoskin data, although in a lesser degree. For example, a few cells of the PEP type are separated from the other cells of the same type, but in the scRMD imputed data, other than one cell, all PEP cells are clustered together. Within-group and between-group distance ratio also shows that imputation of scRMD can make the single cells in the same group closer to each other (Table S3).

### 3.5 Imputation for Ultra-high-throughput scRNA-seq data

Recently developed ultra-high-throughput scRNA-seq methods such as Drop-seq (Macosko *et al.*, 2015) inDrop (Zilionis *et al.*, 2017) and 10X Genomics Chromium (Zheng *et al.*, 2017) can profile thousands of single cells simultaneously. These methods usually use unique molecular identifiers (UMI) to barcode single cells and single RNA moleculars. UMI counts are used for gene expression measurements. These UMI based ultra-high-throughput methods generally have a higher percentage of zeros than low throughput methods. In this subsection, we evaluate the performance of scRMD on these ultra-high-throughput scRNA-seq. We consider three data sets, the Alles data (Alles *et al.*, 2017), the Karaiskos data (Karaiskos *et al.*, 2017) and the Hrvatin data (Hrvatin *et al.*, 2018). Both Alles data and Karaiskos data were generated by the Drop-seq platform and the Hrvatin data was generated by the inDrop platform. The Alles data composes of live human/mouse single cells and chemically fixed human/mouse single cells. The Karaiskos data consists of single Drosophila embryo cells at different regions of the Drosophila embryo. The Hrvatin data is mouse visual cortex single cell data. After preprocessing, we imputed the data by scRMD, MAGIC and scImpute. Figure 6 showed the tSNE visualization of the raw and scRMD-imputed Karaiskos and Hrvatin data. The tSNE visualization of the Alles data and the imputed data by MAGIC and scImpute is shown in Figure S5 and S6. For the Karaiskos data, scRMD clearly demonstrates better visualization than other methods. Compared with the raw data, after scRMD imputation, the region 5 single cells are now tightly clustered together. Furthermore, the region 3 and 4 single cells are more tightly clustered together than the raw data. For the Hrvatin data, scRMD also performs better in visualization of tSNE. Single cells of different classes are further separated and of the same classes become closer together than the raw data. Especially, the ExcL23_1 single cells are largely separated to two clusters in the raw data plot, but they become largely one cluster in the scRMD plot. In comparison, the ExcL23_1 single cells remain to be largely two clusters in the MAGIC and scImpute plots. Overall, all tSNE plots of the Alles data separate single cells from different single cell groups. Interestingly, in the scRMD plot, similar types of human and mice cells are always largely next to each other. For example, the live cells of human and mice are both at the bottom of the plot. This is not the case for the other plots. To further compare the with-group and between-group distance ratios, we find that scRMD gives the smallest ratio value in these data sets. For example, the ratio of scRMD-imputed Karaiskos and Hrvatin data are only 0.55 and 0.38, respectively, much smaller than other methods (Table S4).

**Fig. 6.**
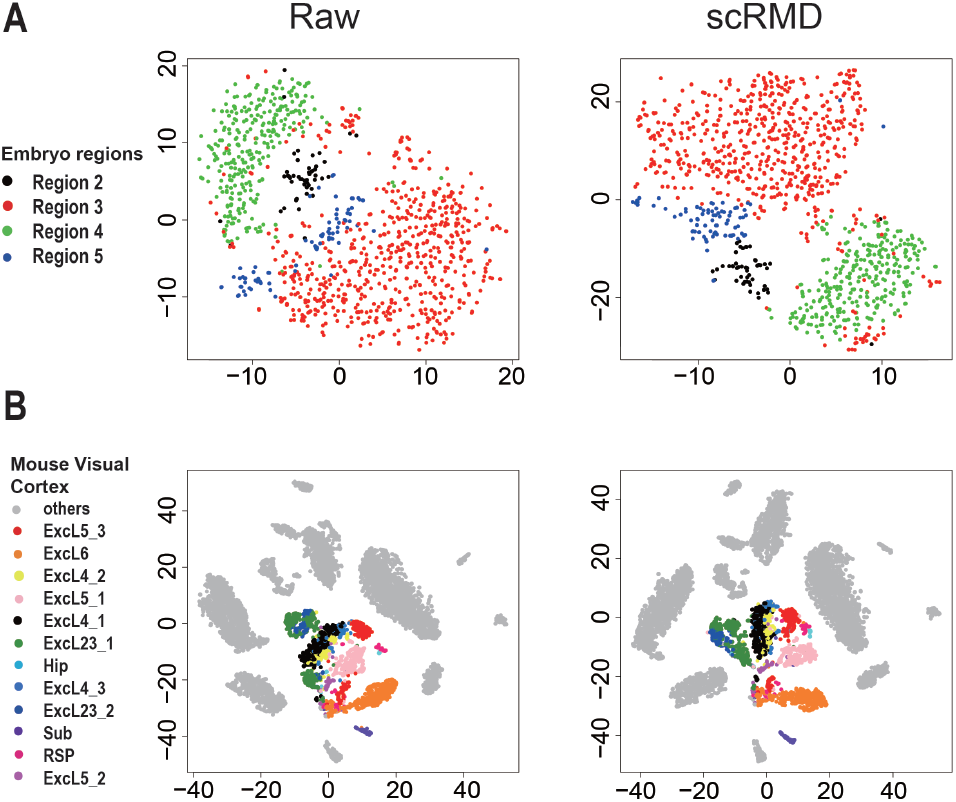
The tSNE visualization of the raw and scRMD imputed data. (A) The Karaiskos data set. (B) The Hrvatin data.

## 4 Discussion

Dropout is well-known to be a major issue in scRNA-seq data. To address this dropout problem, we develop an scRNA-seq data imputation method called scRMD. scRMD only requires two minimal assumptions, the low-rankness assumption and the sparsity assumption. The low-rankness assumption can be naturally satisfied since scRNA-seq data is often composed of cells from several cell types. The sparsity assumption is also reasonable as long as the sequencing depth is not too shallow. The objective function of scRMD has two tuning parameters corresponding to the two penalties. We use random matrix theory to guide the selection of these two parameters and thus avoid the computationally expensive parameter tuning procedure. Extensive data analyses show that this choice of the tuning parameters renders good performance. In the current scRMD, we set the candidate dropout set as the set of genes with expression less than 0.05 quantile of the nonzero expression. Although we find that this choice of candidate dropout set works well for our simulation and real data, the performance of scRMD may be further improved if we introduce another step to select a best threshold *c*. This may be achieved by bootstrapping and measuring the stability of the imputed values of the bootstrapped data. We expect that scRMD can find its applications in many different scRNA-seq studies.

## Acknowledgements

The authors gratefully acknowledge Peter V. Kharchenko for his helpful suggestions.

